# Regulation of pDC Fate Determination by Histone Deacetylase 3

**DOI:** 10.1101/2022.06.08.494949

**Authors:** Yijun Zhang, Zhimin He, Wenlong Lai, Xiangyi Shen, Tao Wu, Jiaoyan Lv, Li Wu

**Affiliations:** Institute for Immunology, Tsinghua-Peking Center for Life Sciences, School of Medicine, Tsinghua University, Beijing 100084, China; Beijing Key Laboratory for Immunological Research on Chronic Diseases, Beijing 100084, China

## Abstract

Dendritic cells (DCs), the key antigen-presenting cells, are primary regulators of immune responses. Transcriptional regulation of DC development had been one of the major research interests in DC biology, however, the epigenetic regulatory mechanisms during DC development remains unclear. Here, we report that *Histone deacetylase 3* (*Hdac3*), an important epigenetic regulator, is highly expressed in pDCs, and its deficiency profoundly impaired the development of pDCs. Significant disturbance of homeostasis of hematopoietic progenitors was also observed in HDAC3-deficient mice, manifested by altered cell numbers of these progenitors and defective differentiation potentials for pDCs. Using the *in vitro* Flt3L supplemented DC culture system, we further demonstrated that HDAC3 was required for the differentiation of pDCs from progenitors at all developmental stages. Mechanistically, HDAC3 deficiency resulted in enhanced expression of cDC1-associated genes, owing to markedly elevated H3K27 acetylation (H3K27ac) at these gene sites in BM pDCs. In contrast, the expression of pDC-associated genes was significantly downregulated, leading to defective pDC differentiation.

**Summary:** This work reveals for the first time that HDAC3 is required for the development of pDCs and maintenance of homeostasis of hematopoietic progenitors with DC differentiation potential. Mechanistically, HDAC3 promotes pDC development by repressing the expression of cDC1 associated genes in a deacetylase dependent manner.

## Introduction

Dendritic cells (DCs) are essential regulators of immune responses. Major DC subsets include conventional dendritic cells (cDCs) and plasmacytoid dendritic cells (pDCs). cDCs are professional antigen-presenting cells that are critical for initiation of antigen specific adaptive immunity and induction of immune tolerance. pDCs are the major producers of large amounts of type I interferon (IFN) during viral infection(Reizis, 2019). In the past decades, the knowledge on the cytokines, transcription factors, and progenitors involved in DC development had been obtained(Nutt and Chopin, 2020; Anderson et al., 2021). However, the role of epigenetic regulation in DC lineage determination and differentiation remained elusive.

The ultimate origin of DCs is the hematopoietic stem cells (HSC), which give rise to multipotent progenitors (MPPs) with potential to populate entire hematopoietic system. MPPs produce common lymphoid progenitors (CLPs) and common myeloid progenitors (CMPs), which give rise to lymphoid and myeloid lineage cells respectively. Both CLPs and CMPs have the potential to differentiate into DCs(Sathe et al., 2013; Wu et al., 2001; D’Amico and Wu, 2003; Chicha et al., 2004; Manz et al., 2001). CMPs further develop into common DC precursors (CDPs) with restricted potential for DC lineage differentiation(Naik et al., 2007; Onai et al., 2007), CDPs differentiate into pDC and pre-cDCs in the bone marrow(Liu and Nussenzweig, 2010; Puhr et al., 2015). The CD115-CDP subsets predominantly produce pDC, while CD115^+^CDP subsets produce more cDCs(Onai et al., 2013). The Siglec H^+^Ly6D^+^ subset of LPs (CLPs) exhibit a specific potential to differentiate towards pDC(Rodrigues et al., 2018; Dress et al., 2019).

Several transcription factors and key regulators have been identified to play important roles in pDC development. Transcription factor TCF4 (E2-2) is highly expressed in pDCs(Nagasawa et al., 2008). Deletion of TCF4 disrupts the development and function of pDCs(Cisse et al., 2008; Ghosh et al., 2010). The development of pDC also require repressed expression of ID2, as well as PU.1, MTG16, ZEB2 and BCL11A (Schlitzer et al., 2011; Carotta et al., 2010; Ghosh et al., 2014; Scott et al., 2016; Wu et al., 2016; Ippolito et al., 2014; Wu et al., 2013; Luc et al., 2016; Nagasawa et al., 2008).

Histone deacetylase 3 (HDAC3) is a key epigenetic regulator orchestrating histone modification and chromatin remodeling(Emmett and Lazar, 2019). Previous studies showed that HDAC3 regulates stem cell differentiation(Summers et al., 2013), T and B lymphocyte development(Stengel et al., 2017, 2019, 2015; Philips et al., 2016; Hsu et al., 2015), and the function of macrophages(Chen et al., 2012; Nguyen et al., 2020). Generation of DCs *in vitro* in the presence of pan-HDAC inhibitors revealed that HDACs were required for establishing a DC gene network(Chauvistré et al., 2014). However, until now which specific HDAC plays the major role in the development of DCs remains to be elucidated.

In this study, we observed a higher expression of *Hdac3* in pDCs compared to that of cDCs. The development of pDCs was impaired in HDAC3-deficient mice, mainly due to the alterations in pDC differentiation potential of the HDAC3-deficient DC progenitors. Mechanistically, HDAC3 was required for repressing the H3K27ac level at cDC1-associated gene loci such as *Zfp366, Batf3* and *Zbtb46* etc., thereby repressing the expression of cDC1-associated genes and promoting the differentiation towards pDCs. Together, our study revealed a crucial role of HDAC3 in regulating the development of pDCs from multiple hematopoietic progenitors with DC differentiation potential, and the histone deacetylase activity of HDAC3 is required for this process. Our findings provide novel insights into the epigenetic regulatory mechanisms for pDC development.

## Results

### HDAC3 deficiency resulted in defective development of pDCs

Genome-wide expression dataset ImmGen reveals that among class I HDACs, *Hdac1, Hdac2* and *Hdac3* but not *Hdac8* are highly expressed by most immune cells. *Hdac3* is mostly expressed in hematopoietic progenitors such as HSCs and CLP, and pDCs in spleen (**Figure 1- figure supplement 1A**). Consistent with these data, *Hdac3* expression in sorted DC subsets analyzed by quantitative real-time PCR (qRT-PCR) showed that pDC expressed significantly higher levels of *Hdac3* than cDCs (**Figure 1- figure supplement 1B**). Thus, we speculated that *Hdac3* might play important roles in pDC development or function.

**Figure 1.**
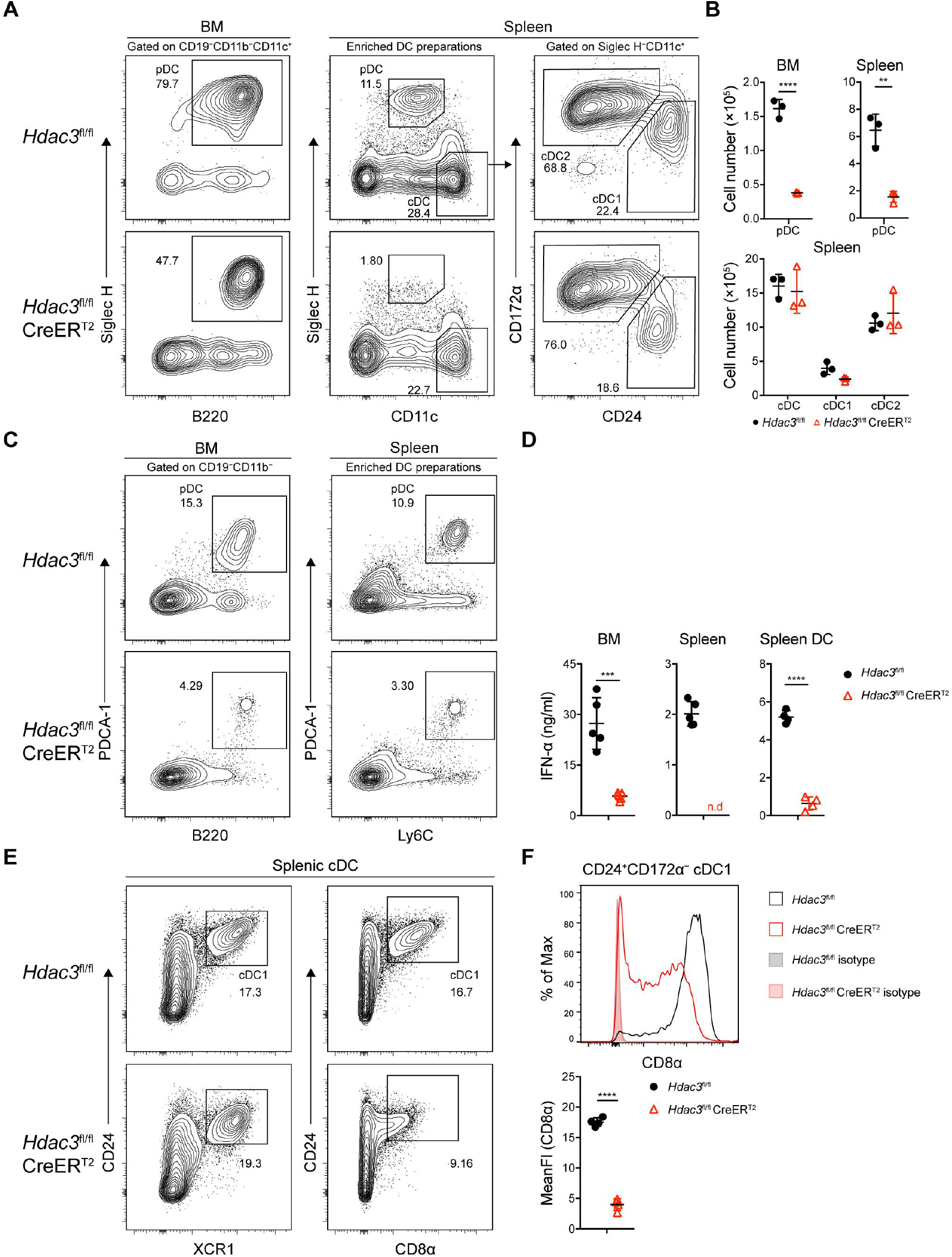
Impaired pDC development in *Hdac3*-deficient mice. **(A-B)** Representative flow cytometry profiles of BM pDCs and splenic DC subsets (A) and the absolute numbers of each DC subset of *Hdac3*^fl/fl^ and *Hdac3*^fl/fl^ CreER^T2^ mice (B). **(C)** The pDC populations in BM and spleen were defined using a combination of indicated markers. Shown are staining profiles of gated CD11b^−^CD19^−^ cells from the BM and DC enrichment preparations from the spleens. **(D)** IFN-α production by *Hdac3*^fl/fl^ or *Hdac3*^fl/fl^ CreER^T2^ cells *in vitro*. Total BM cells, splenocytes or purified DC preparations from the spleens were stimulated with CpG ODN 2216, and IFN-α in the supernatants was measured after 18 hours by ELISA. Cells were harvested from 4-5 mice each group. **(E)** The cDC1 populations in spleen were defined using a combination of indicated markers. Staining profiles of gated Siglec H^−^CD11c^+^ cells from the spleen are shown. **(F)** The expression and mean fluorescent intensities (MFIs) of CD8α in splenic CD24^+^CD172α^−^ cDC1s from *Hdac3*^fl/fl^ and *Hdac3*^fl/fl^ CreER^T2^ mice. Results are from one experiment representative of 3 independent experiments with 3 mice per group. Data are shown as mean ± SD. * *p* < 0.05; ** *p* < 0.01; *** *p* < 0.001; **** *p* < 0.0001, by two-tailed Student’s *t*-test.

To test the role of HDAC3 in pDC development, *Hdac3*^fl/fl^ mice were crossed with CreER^T2^ mice to generate HDAC3 conditional knockout mice (**Figure 1- figure supplement 2A**). *Hdac3*^fl/fl^ CreER^T2^ mice and *Hdac3*^fl/fl^ controls mice were treated with Tamoxifen (intraperitoneal injection, i.p.) for 5 consecutive days and DC subsets were analyzed 7 days after the last tamoxifen administration (**Figure 1- figure supplement 2B**). HDAC3 could be efficiently deleted in *Hdac3*^fl/fl^ CreER^T2^ mice (**Figure 1- figure supplement 2C and 2D, Figure 1- figure supplement 2D source data 1**). Murine pDCs express low level CD11c, high level B220, CD45RA, Ly6C, and can be defined by specific markers such as Siglec H and PDCA-1(Anderson et al., 2021). Compared to *Hdac3*^fl/fl^ mice, *Hdac3*^fl/fl^ CreER^T2^ mice showed substantially reduced pDCs and the absolute number decreased about 4-fold in BM and spleens (**Figure. 1A and B**), whereas the absolute numbers of CD11c^hi^ cDCs, CD24^+^CD172α^−^cDC1s and CD24^−^CD172α^+^ cDC2s were comparable (**Figure. 1A and B**). To further confirm the number of pDC was reduced additional pDC surface markers PDCA-1 and Ly6C were also applied to identify pDC population, and all showed consistent reduction in pDC numbers in BM and spleen of *Hdac3*^fl/fl^ CreER^T2^ mice (**Figure. 1C**). pDCs are the major producers of type-I IFN, consistent with the reduction of pDCs in BM and spleen of *Hdac3*^fl/fl^ CreER^T2^ mice, the production of IFN-α by *Hdac3*^fl/fl^ CreER^T2^ BM, splenocytes and splenic DCs were also decreased in response to type A CpG ODN stimulation *in vitro* (**Figure. 1D**). Splenic cDC1 can also be identified using cell surface markers including XCR1 and CD8α. As shown in **Figure. 1E and F**, in accordance with cDC1 identified by CD24, the percentage of splenic cDC1 marked by XCR1 showed little difference in *Hdac3*^fl/fl^ CreER^T2^ mice compared with that of *Hdac3*^fl/fl^ mice. However, the expression of CD8α significantly decreased in HDAC3-deficient splenic cDC1 cells. Taken together, these results demonstrated the requirement of HDAC3 in pDC development *in vivo*.

### HDAC3 regulated pDC development in a cell-intrinsic manner

To test whether the function of HDAC3 in pDC development is cell-intrinsic, competitive BM chimeric mice were generated with BM cells from CD45.2^+^ *Hdac3*^fl/fl^ or *Hdac3*^fl/fl^ CreER^T2^ mice mixed with CD45.1^+^ WT BM cells at 1:1 ratio. Eight weeks later the chimeric mice were treated with Tamoxifen and the CD45.2^+^ donor derived DC subsets were analyzed by flow cytometry. The percentage of *Hdac3*^fl/fl^ CreER^T2^ BM derived CD45.2^+^ cells were significantly lower than those derived from *Hdac3*^fl/fl^ BM, indicating a competitive disadvantage of the HDAC3-deficient BM cells (**Figure. 2A**). The percentage of CD45.2^+^ cells derived from *Hdac3*^fl/fl^ CreER^T2^ donors only represented about 20% in BM and 30% in spleen of the chimeric mice (**Figure. 2B**). Furthermore, the frequency of *Hdac3*^fl/fl^ CreER^T2^ derived BM and splenic pDCs among CD45.2^+^ cells also decreased markedly, whereas the frequency of splenic cDCs increased due to the reduction of pDCs (**Figure. 2, A-C**). Moreover, when the BM cells were cultured *in vitro* for 5, 7 and 9 days in the presence of FLT3L and 4-hydroxytamoxifen (4-OHT) to induce *Hdac3* deletion in *Hdac3*^fl/fl^ CreER^T2^ BM cells, impaired pDC development from *Hdac3* deleted BM cells was also observed, in line with the results from *in vivo* study (**Figure. 2D and E**), suggesting that HDAC3 was required for pDC development in a cell intrinsic manner.

**Figure 2.**
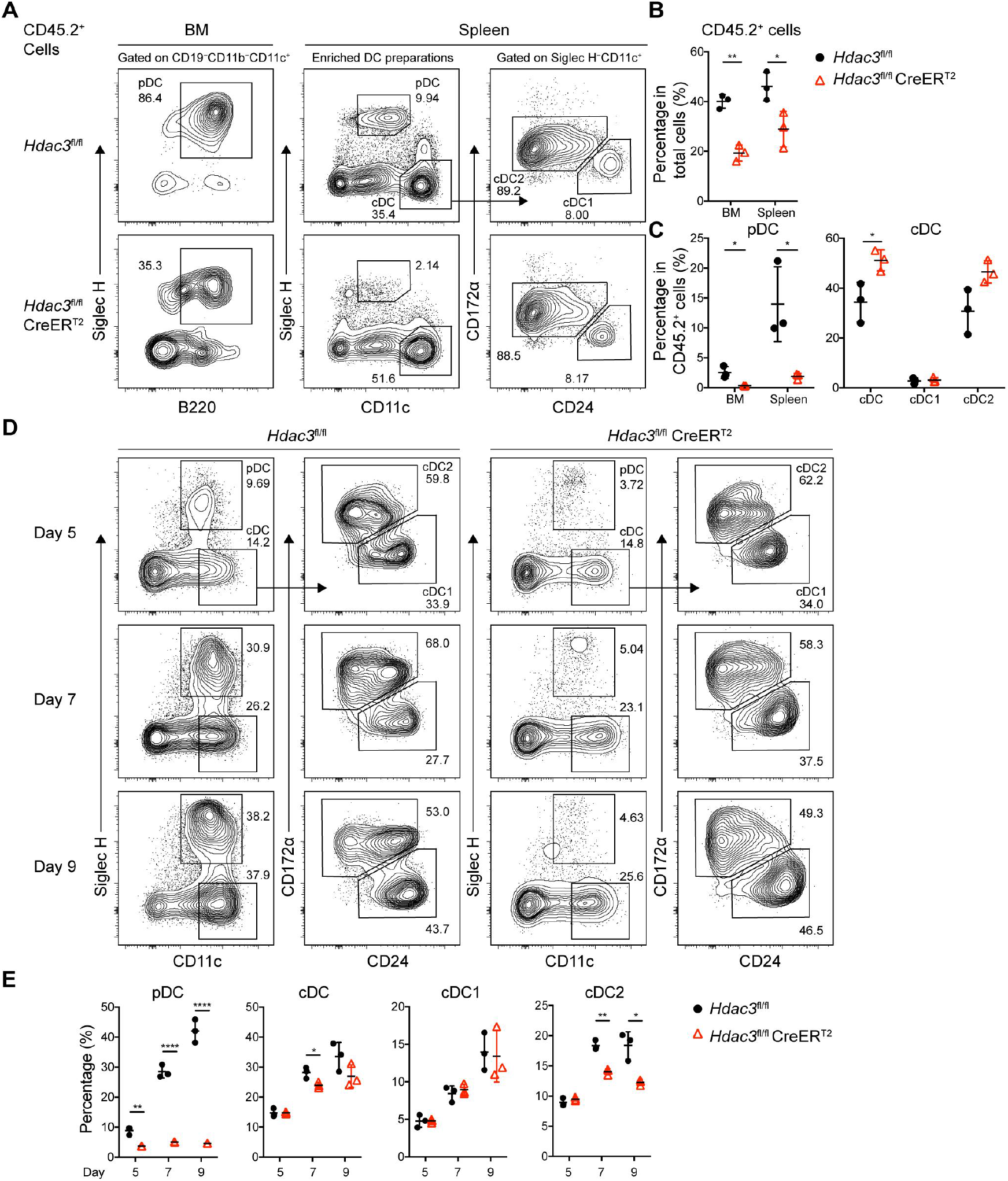
HDAC3 regulated pDC development in a cell-intrinsic manner. Lethally irradiated CD45.1 WT mice were reconstituted with a mixture of CD45.1 WT BM and BM from CD45.2 *Hdac3*^fl/fl^ or *Hdac3*^fl/fl^ CreER^T2^ mice at 1:1 ratio. 8 weeks after reconstitution, *Hdac3* deletion was induced by tamoxifen. **(A-C)** Results shown are staining profiles of gated CD11b^−^CD19^−^CD11c^+^ cells from the BM and enriched DC preparations from the spleens. Representative flow cytometry profiles of CD45.2^+^ donor derived BM pDCs and splenic DC subsets (A), the percentage of CD45.2^+^ cells in BM and spleen (B), and the percentage of each DC subset among CD45.2^+^ cells (C) in the BM chimeric mice. Results are from one experiment representative of 3 independent experiments with 3 animals per group. **(D-E)** Total BM cells were plated at 1.5×10^6^ cells/ml in the presence of 200ng/ml FLT3L and 1μM 4-Hydroxytamoxifen (4OH-T). Results shown representative flow cytometry profiles (D) and the percentage of DC subsets (E) of FLT3L stimulated BM cultures on day 5, 7 and 9. Results are from one experiment representative of 3 independent experiments with 3 animals per group. Data are shown as mean ± SD. * *p* < 0.05; ** *p* < 0.01; *** *p* < 0.001; **** *p* < 0.0001, by two-tailed Student’s *t*-test.

### HDAC3 deletion led to disturbed homeostasis of early hematopoietic progenitors

To further investigate whether the absence of pDC in HDAC3^-^deficient mice was mainly due to decreased numbers of hematopoietic progenitors or resulted from inability of HDAC3^-^deficient progenitors to differentiate into pDCs, we first examined the composition of BM progenitor populations. BM Lin^−^ cells from *Hdac3*^fl/fl^ control or *Hdac3*^fl/fl^ CreER^T2^ mice after Tamoxifen treatment were analyzed for each progenitor populations. Compared to control mice, the Lin^−^Sca-1^+^CD117^+^ (LSK) cells were increased significantly in *Hdac3*^fl/fl^ CreER^T2^ mice. Further analysis of LSK subpopulations, namely the CD34^−^Flt3^−^ long-term HSC (LT-HSC), CD34^+^Flt3^−^ short-term HSC (ST-HSC) and Flt3^+^CD34^+^ lymphocyte-primed multipotent precursors (LMPP), showed increased numbers of all three subpopulations (**Figure. 3A and B**). Meanwhile, HDAC3 deficiency led to markedly decreased number of Sca-1^−^CD117^+^CD16/32^int^CD34^+^ CMP, while increased number of Sca-1^−^CD117^+^CD16/32^+^CD34^+^ granulocyte-macrophage progenitor (GMP), but comparable number of Sca-1^−^CD117^+^CD16/32^−^CD34^−^ megakaryocyte-erythroid progenitor (MEP) (**Figure. 3, A-B**).The major pDC precursor populations are the CD117^int^Flt3^+^CD127^+^ CLPs and the CD117^int^Flt3^+^CD127^−^CD11c^−^ CDPs which include both CD115^+^ and CD115^−^ subsets, whereas cDCs are mainly produced by CD115^+^CDP(Rodrigues et al., 2018; Dress et al., 2019; Onai et al., 2007, 2013). HDAC3-deficient mice showed a significant reduction in the numbers of CD115^+^CDP and CLP, while unexpectedly the number of CD115^−^CDP cells increased significantly (**Figure. 3, C-F**). A schematic diagram summarizing the changes in numbers of hematopoietic progenitors with DC differentiation potential in HDAC3-deficient mice was shown in **Figure. 3G**. Apart from the increased number of LSKs and CD115^−^CDPs, the number of all the other progenitors with DC differentiation potential decreased significantly in HDAC3-deficient mice.

**Figure 3.**
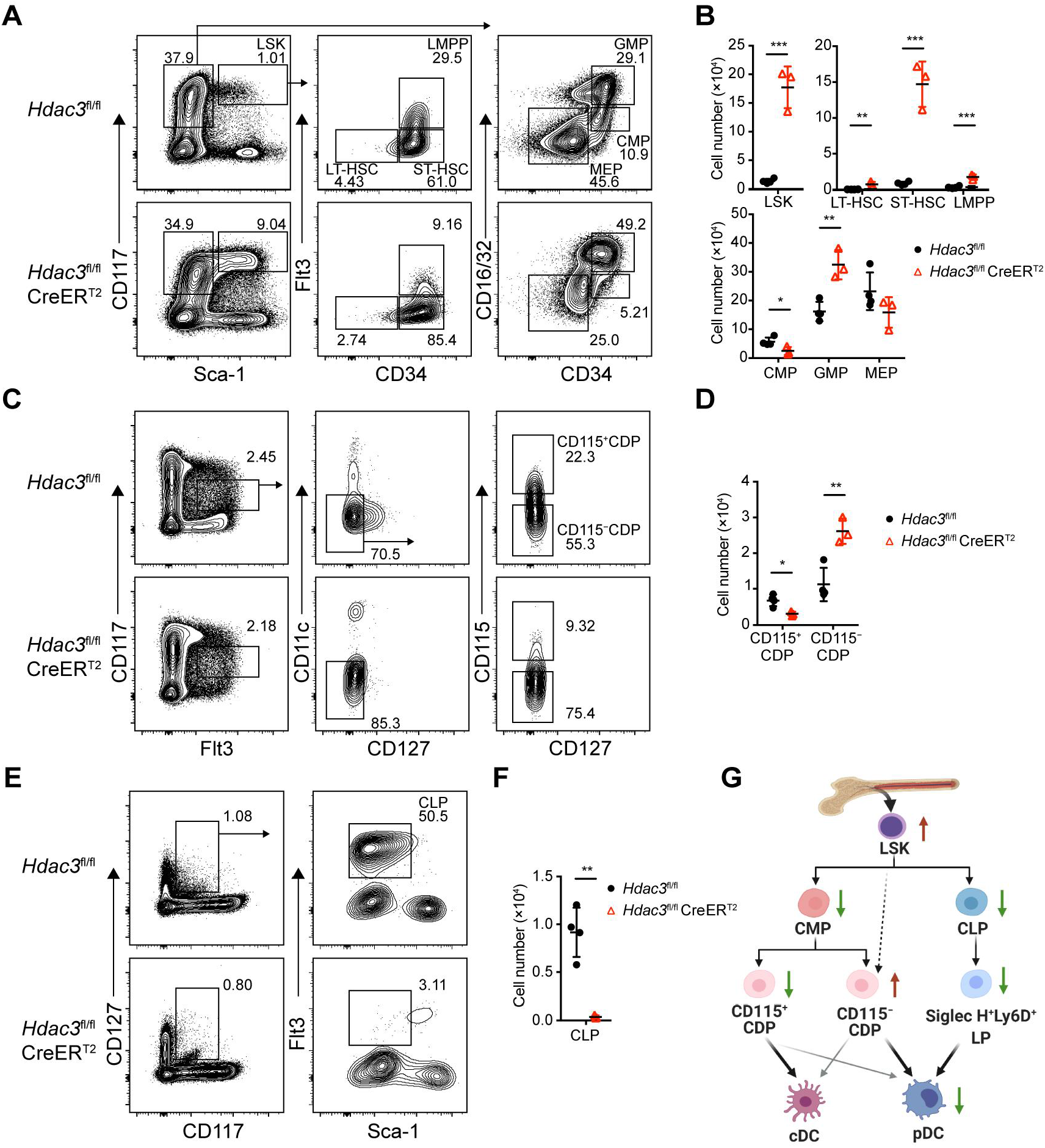
HDAC3 deficiency resulted in altered numbers of DC progenitor populations. **(A-B)** Representative flow cytometry profiles (A) and the absolute cell number (B) of indicated progenitor populations, LSK, CMP, GMP, MEP within *Hdac3*^fl/fl^ or *Hdac3*^fl/fl^ CreER^T2^ BM Lin^−^ cells. **(C-D)** Representative flow cytometry profiles (C) and the absolute cell number (D) of indicated CDP populations. **(E-F)** Representative flow cytometry profiles (E) and the absolute cell number (F) of CLP. **(G)** A schematic diagram summarizing the changes in numbers of progenitors with DC differentiation potential in HDAC3-deficient mice. Red arrow: cell number increase, green arrow: cell number reduction. Diagram was created with BioRender.com. Results are from one experiment representative of at least 2 independent experiments with 3-4 animals per group. Data are shown as mean ± SD. * *p* < 0.05; ** *p* < 0.01; *** *p* < 0.001; **** *p* < 0.0001, by two-tailed Student’s *t*-test.

It was reported that HDAC3 was required for the passage of HSCs through regulating S phase and the formation of early lymphoid progenitors(Summers et al., 2013). We then measured the DNA synthesis in the DC progenitor cells during proliferation using BrdU-incorporation assay. Mice treated with Tamoxifen and then injected i.p. with BrdU 24 hours before analysis. Compared with that of control cells, HDAC3-deficiency led to an increased percentage of BrdU-incorporated LSKs, but decreased proportion of BrdU-incorporated CLPs, and comparable ratios of BrdU-incorporated CD115^−^CDPs and CD115^+^CDPs respectively (**Figure 3 - Figure supplement 1 A and B**). These results suggested that the disturbed homeostasis of progenitor cells caused by HDAC3 deficiency might mainly due to their altered proliferation rates.

Taken together, the disturbed homeostasis of BM progenitor cells might contribute to the defective pDC development in HDAC3-deficient mice. It remained unclear whether the differentiation potentials for pDCs of these progenitors were also affected by HDAC3 deficiency.

### HDAC3 deficiency abrogated pDC differentiation potential of LSKs and CD115^−^CDPs

As described above, HDAC3-deficient mice exhibited increased numbers of LSK cells and CD115^−^CDPs. To examine whether these increased progenitors could still give rise to pDCs, LSK cells and CD115^−^CDPs were purified from the BM of Tamoxifen treated CD45.2 *Hdac3*^fl/fl^ or *Hdac3*^fl/fl^ CreER^T2^ mice and then transplanted into lethally radiated CD45.1 recipient mice together with CD45.1 total BM competitors (**Figure. 4A**). HDAC3-deficient LSKs and CD115^−^CDPs showed significant competitive disadvantage, revealed by the very low percentage of CD45.2^+^ cells in chimeric mice generated with HDAC3-deficient progenitors compared with that of *Hdac3*^fl/fl^ progenitors (**Figure. 4B and C**). Furthermore, a decreased frequency of pDCs derived from HDAC3-deficient CD115^−^CDP was detected in both BM and spleen, whereas the frequencies of cDC subsets in the spleen were comparable to that derived from control CD115^−^CDP. These results indicated that HDAC3 deficiency in CD115^−^CDPs resulted in severe defect in pDC, but not in cDC development (**Figure. 4D and E**). To further explore the regulatory role of HDAC3 in these progenitors, we compared gene expression profiles of LMPPs (the Flt3^+^CD34^+^ subset of LSKs) and CD115^−^CDPs from Tamoxifen treated *Hdac3*^fl/fl^ and *Hdac3*^fl/fl^ CreER^T2^ mice by RNA-seq analysis. HDAC3 deficiency resulted in down-regulation of 2533 and up-regulation of 206 genes with annotations in LMPPs (**Figure. 4F**). KEGG pathway analysis of differentially expressed genes revealed a significant enrichment of genes for hematopoietic lineage pathway (**Figure. 4G**), suggesting that HDAC3 may play important roles in hematopoietic linage cell differentiation. Interestingly, the HDAC3-deficient CD115^−^CDPs showed down regulated expression of genes that are highly expressed in pDC, such as *Siglech, Ly6c2, Ly6d, Cd209a* and *Cox6a2* (**Figure. 4H**). In addition, myeloid-related genes such as *Mpo, Ctsg* and *Ms4a3* were up-regulated, but pDC-associated genes such *Siglech, Ly6d* and *Ly6c2* were down-regulated in both LMPP and CD115^−^CDPs (**Figure. 4I**). These results indicated that HDAC3 is crucial for the expression of pDC-associated genes in those progenitors. Overall, these results indicated that HDAC3-deficient LSKs and CD115^−^CDPs were defective to differentiate into pDCs even though with increased cell numbers.

**Figure 4.**
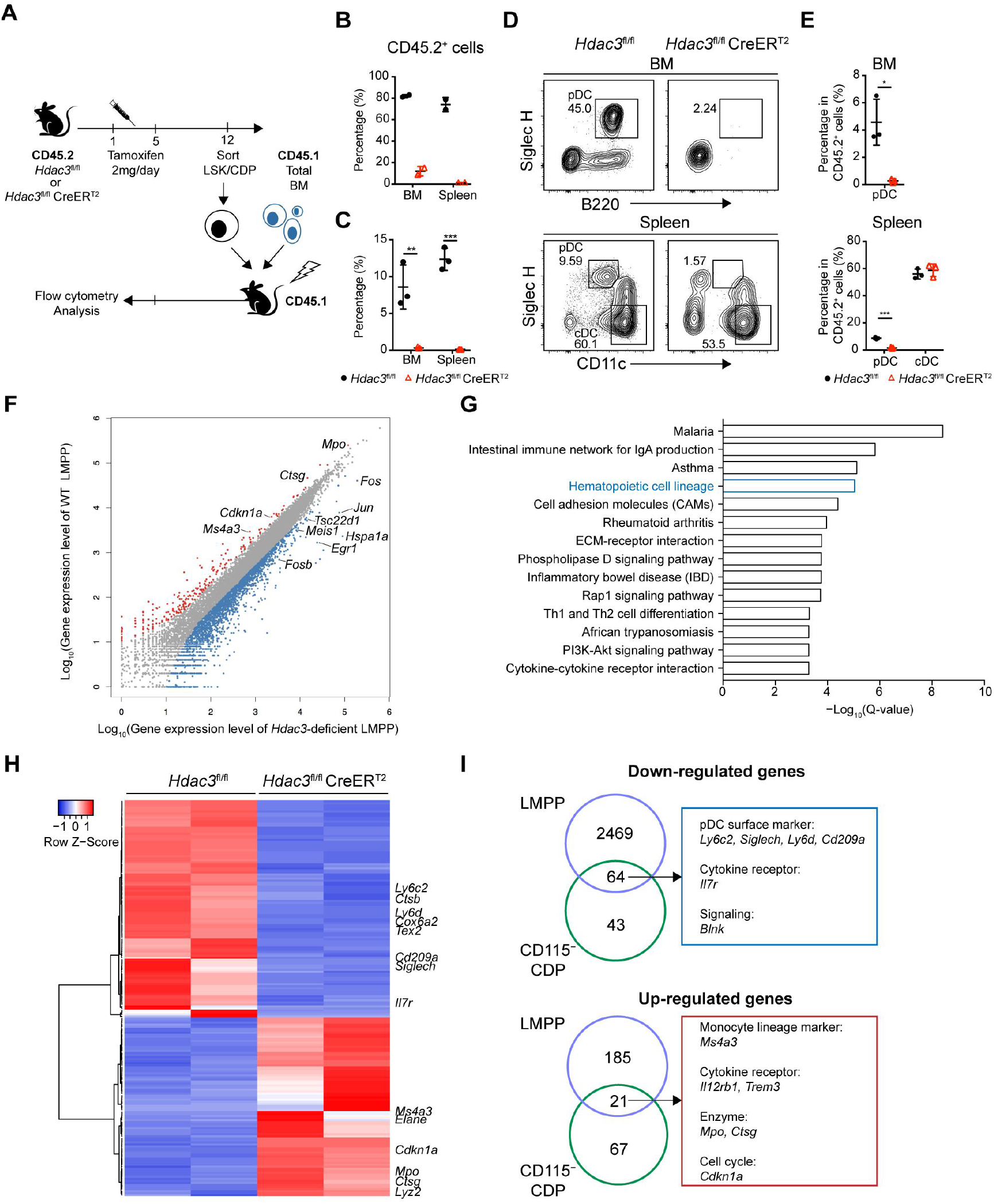
HDAC3 regulated pDC development by modulating the expression of pDC signature genes. **(A)** Schematic diagram of progenitor transplantation experiment. HSCs and CD115^−^CDPs were purified from the BM of *Hdac3*^fl/fl^ and *Hdac3*^fl/fl^ CreER^T2^ mice after tamoxifen treatment, and then transplanted together with CD45.1 BM competitors into lethally irradiated CD45.1 WT mice. Three weeks (LSK) or 10 days (CD115^−^CDPs) post reconstitution, BM and spleen cells from BM chimeric mice were analyzed for pDC and cDC repopulation. **(B-C)** The percentage of CD45.2^+^ cells in BM chimeric mice repopulated with LSK (B) or CD115^−^CDP (C). **(D-E)** Representative flow cytometry profiles of BM pDCs and splenic DC subsets in CD45.2^+^ cells (D) and the percentage of DC subsets among CD45.2^+^ cells in the BM and spleen (E) of BM chimeric mice reconstituted with CD115^−^CDPs. Results are from one experiment representative of at least three independent experiments with 2-3 animals per group. Data are shown as mean ± SD. * *p* < 0.05; ** *p* < 0.01; *** *p* < 0.001; **** *p* < 0.0001, by two-tailed Student’s *t*-test. **(F)** Scatter plot shows the differentially expressed genes (DEGs) (Fold Change > 2, q-value ≤ 0.001), which were up-regulated (red) and down-regulated (blue) in *Hdac3*-deficient LMPPs compared with WT LMPPs. Duplicate samples for each genotype were analyzed. **(G)** KEGG pathway analysis of DEGs in *Hdac3*-deficient LMPPs compared with WT LMPPs. Shown are pathways with Q-value ≤ 0.001. **(H)** Heatmap shows the DEGs (Fold Change > 2, q-value ≤ 0.05) between *Hdac3*-deficient and WT CD115^−^CDPs. Duplicate samples for each genotype were analyzed. **(I)** Common genes down-regulated or up-regulated in *Hdac3*-deficient LMPPs and CD115^−^CDPs compared with those from WT LMPP and CD115^−^CDPs.

### HDAC3 was required for pDC development from multiple early hematopoietic progenitors

To further clarify whether HDAC3 is required for pDC development from hematopoietic progenitors at different developmental stages, we deleted *Hdac3* in BM cells or different progenitors by applying 4OH-T into FLT3L stimulated cultures with BM cells from *Hdac3*^fl/fl^ or *Hdac3*^fl/fl^ CreER^T2^ mice on day 0, 2, 4 or 6, and analyzed the generation of DC subsets on day 8. Deletion of *Hdac3* at the beginning (day 0) of the BM culture totally prohibited pDC development. Whereas some pDCs could still be generated from in the cultures when *Hdac3* was deleted at later stages (e.g.day 4 and day 6) (**Figure. 5A and B**). We also analyzed the *Hdac3*^fl/fl^ *Itgax*-Cre mice, in which *Hdac3* was selectively deleted after CDP stage in CD11c^+^ cells (**Figure 5 - Figure supplement 1 A and B, Figure 5- figure supplement 1B source data 1**), to evaluate whether a later stage deletion of *Hdac3* would affect DC development *in vivo*. As shown in **Figure 5 - Figure supplement 1 C and D**, pDCs and cDCs could develop normally in the spleen of *Hdac3*^fl/fl^ *Itgax*-Cre mice. These results indicated a requirement of HDAC3 at early stage of pDC development. As pDCs can develop from Flt3^+^LSKs (LMPPs), Flt3^+^CMPs, CLPs and CDPs, we also tested the requirement of HDAC3 during pDC development from these progenitors. Purified progenitors from *Hdac3*^fl/fl^ or *Hdac3*^fl/fl^ CreER^T2^ mice (CD45.2^+^) were cultured with CD45.1 BM feeders *in vitro* in the presence of FLT3L and 4OH-T. Both pDCs and cDCs generated from HDAC3-deficient LMPPs decreased significantly. Whereas HDAC3-deficient Flt3^+^CMPs, CLPs, CD115^+^CDPs and CD115^-^CDPs all showed profound defective pDC development *in vitro*, with no or slight reduction in cDC development (**Figure. 5C and D**). Taken together, these results demonstrated that HDAC3 was required for pDC development from all early progenitors with DC differentiation potential.

**Figure 5.**
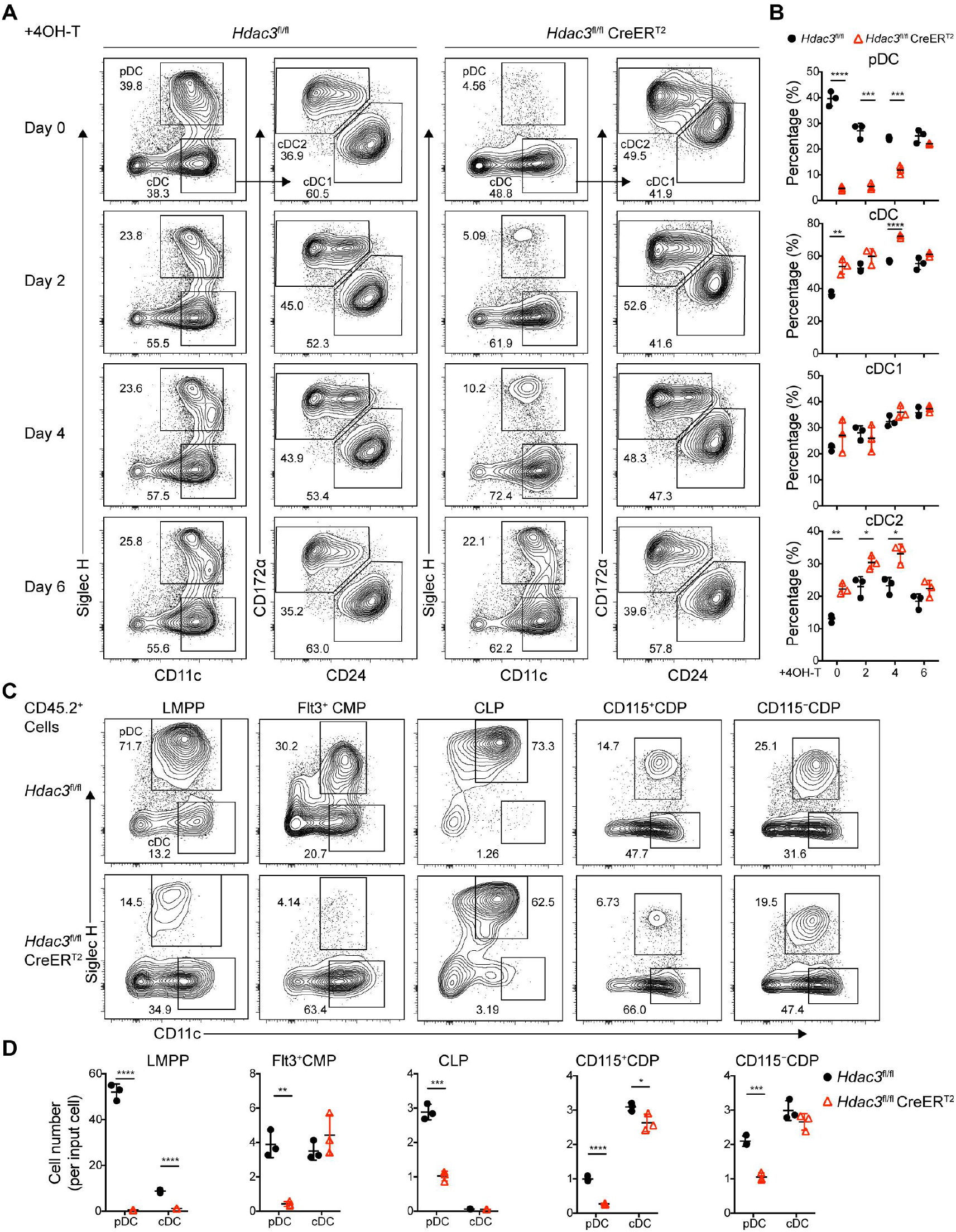
The development of pDCs from BM progenitors required HDAC3. **(A-B)** Total BM cells were plated at 1.5×10^6^ cells/ml in the presence of 200ng/ml FLT3L. 4-Hydroxytamoxifen (4OH-T) was added on day 0, 2, 4 or 6 respectively. Representative flow cytometry profiles (A) and the percentage (B) of each DC subset in FLT3L stimulated BM cultures on day 8. **(C-D)** The progenitor populations LMPP, Flt3^+^CMPs, CLPs, and CDPs, were isolated from BM of *Hdac3*^fl/fl^ or *Hdac3*^fl/fl^ CreER^T2^ mice, and cocultured with CD45.1 BM feeder cells in the presence of FLT3L with 1 μM 4OH-T. Representative flow cytometry profiles (C) and the number of DCs generated in each FLT3L stimulated culture per indicated input progenitor (D). Data are representative of at least two independent experiments with three duplicated wells. Data are shown as mean ± SD. * *p* < 0.05; ** *p* < 0.01; *** *p* < 0.001; **** *p* < 0.0001, by two-tailed Student’s *t*-test.

### Deacetylase activity of HDAC3 was essential for its role in regulating pDC development

HDAC3 has been reported to be a dichotomous transcriptional activator and repressor during the activation of macrophages by LPS, with a non-canonical deacetylase-independent function to activate gene expression(Nguyen et al., 2020). To determine whether the deacetylase activity of HDAC3 is important for its role in regulating pDC development, the HDAC3 selective inhibitor RGFP966(Malvaez et al., 2013) was added into FLT3L stimulated BM culture system. At low concentration, RGFP966 could efficiently inhibit the differentiation of pDCs but not for cDCs. At higher concentration, it exhibited cellular toxicity, evidenced by the dramatic reduction in total cell numbers (**Figure. 6A and B**). This result suggested that HDAC3 deacetylase activity was required preferentially for pDC development. The highly conserved tyrosine residue (Y298) in HDAC3 is located within the active site and is required for HDAC3 catalytic activity. HDAC3 (Y298F) mutant contains a tyrosine-to-phenylalanine substitution at residue 298, which inactivates HDAC3 deacetylase catalytic activity but not the interaction with NCoR/SMRT(Sun et al., 2013). The isolated Lin^−^ BM cells from *Hdac3*^fl/fl^ or *Hdac3*^fl/fl^ CreER^T2^ mice were transduced with empty control vector (GFP), wild-type HDAC3 (WT) or HDAC3 (Y298F) mutant encoding retrovirus, to further confirmed the requirement of HDAC3 deacetylase activity in pDC development (**Figure. 6C**). The amount of HDAC3 (Y298F) protein was comparable to those of HDAC3 (WT) in each group. The development of pDC from HDAC3-deficient Lin^−^ BM cells could be partially rescued by transducing the wild-type HDAC3. However, transduction of the deacetylase-inactivated HDAC3 (Y298F) mutant did not restore pDC development from HDAC3-deficient Lin^−^ BM cells (**Figure. 6D and E**). Together, these results indicated that the deacetylase activity of HDAC3 is essential for its role in regulating the development of pDCs.

**Figure 6.**
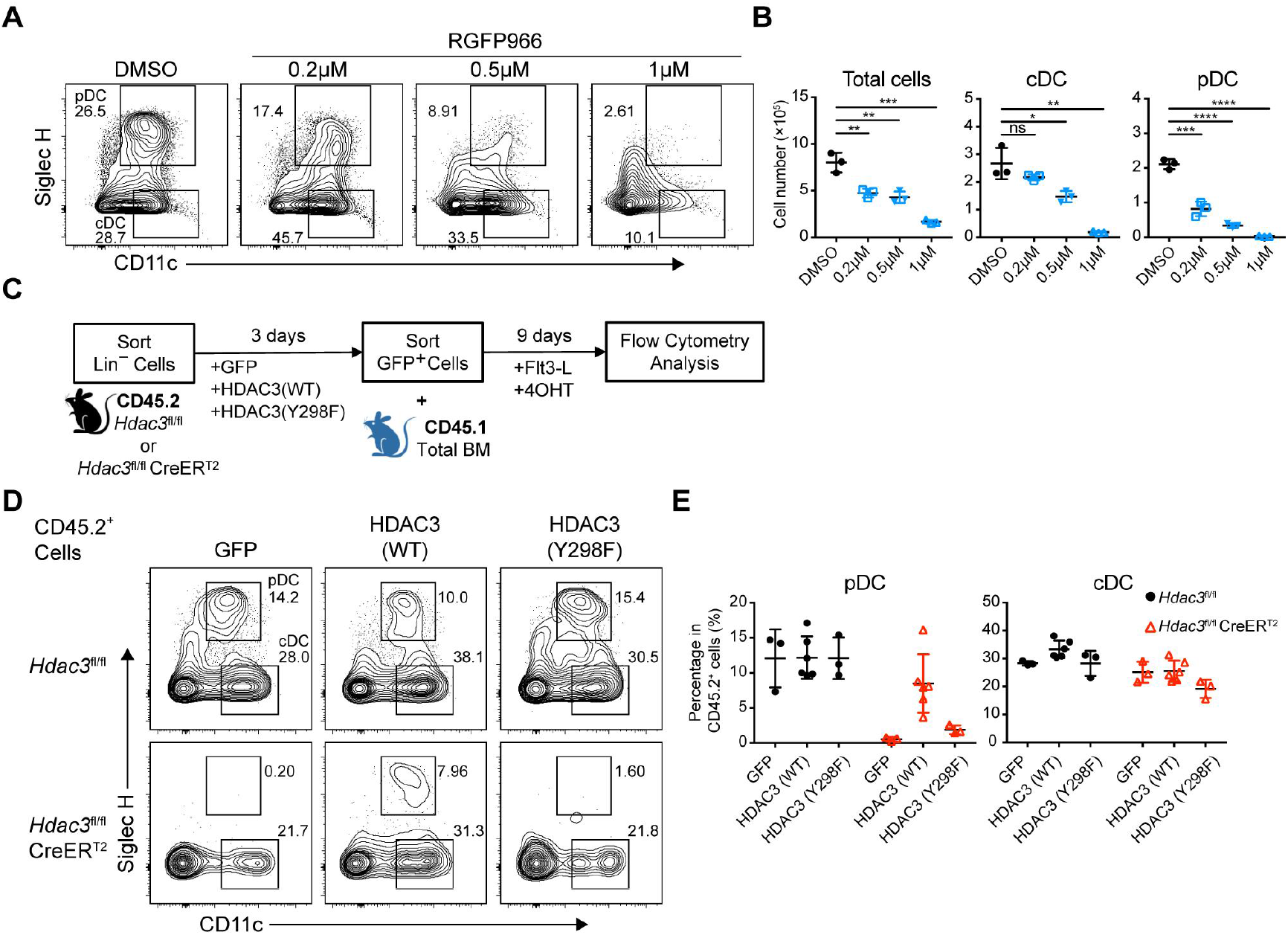
Deacetylase-dependent regulation of pDC development by HDAC3. **(A-B)** Total BM cells were plated at 1.5×10^6^ cells/ml in the presence of 200ng/ml FLT3L and indicated concentration of RGFP966, a selective HDAC3 inhibitor. Representative flow cytometry profiles (A), and the absolute number of total cells, pDC and cDC (B) generated in FLT3L stimulated BM cultures on day 9. Results are representative of at least 2 independent experiments with 3 mice per group. **(C)** BM Lin^−^ cells were transfected with empty (GFP), HDAC3-GFP (WT) and deacetylase-inactivated HDAC3-GFP (indicated as HDAC3(Y298F)) and cultured in the presence of FLT3L for 9 days. **(D-E)** Representative flow cytometry profiles (D), the percentage (E) of DCs generated in FLT3L stimulated cultures from retrovirus-transfected CD45.2^+^ BM Lin^−^ cells. Results are representative of at least two independent experiments with 3-6 duplicated wells per group. Data are shown as mean ± SD. * *p* < 0.05; ** *p* < 0.01; *** *p* < 0.001; **** *p* < 0.0001, by two-tailed Student’s *t*-test.

### HDAC3 regulated H3K27 acetylation level of cDC-associated genes

To explore the mechanism of HDAC3 regulation in pDC development, we compared gene expression profiles of BM pDCs from *Hdac3*^fl/fl^ and *Hdac3*^fl/fl^ CreER^T2^ mice (**Figure. 7, A-D**). Consistent with RNA-seq results obtained from progenitors, HDAC3-deficient pDCs showed down-regulated pDC signature genes such as transcription factor *Tcf4* and pDC surface molecules *Siglech, Ly6c1, Ly6c2* and *Ly6d*. Among up-regulated genes in HDAC3-deficient pDCs, some genes were required in cDC1 development, such as *Spi1*(Dakic et al., 2005; Carotta et al., 2010; Chopin et al., 2019), *Zfp366*(Zhang et al., 2021), *Batf3*(Hildner et al., 2008; Grajales-Reyes et al., 2015) and *Id2*(Jackson et al., 2011; Sivakamasundari et al., 2015). Specific marker of cDC specification *Zbtb46*(Meredith et al., 2012a; b) and the surface molecule highly expressed by cDC1, such as *Clec9a* were also up-regulated in HDAC3-deficient pDCs. These results displayed a cDC-biased transcriptional profile in HDAC3-deficient pDCs.

**Figure 7.**
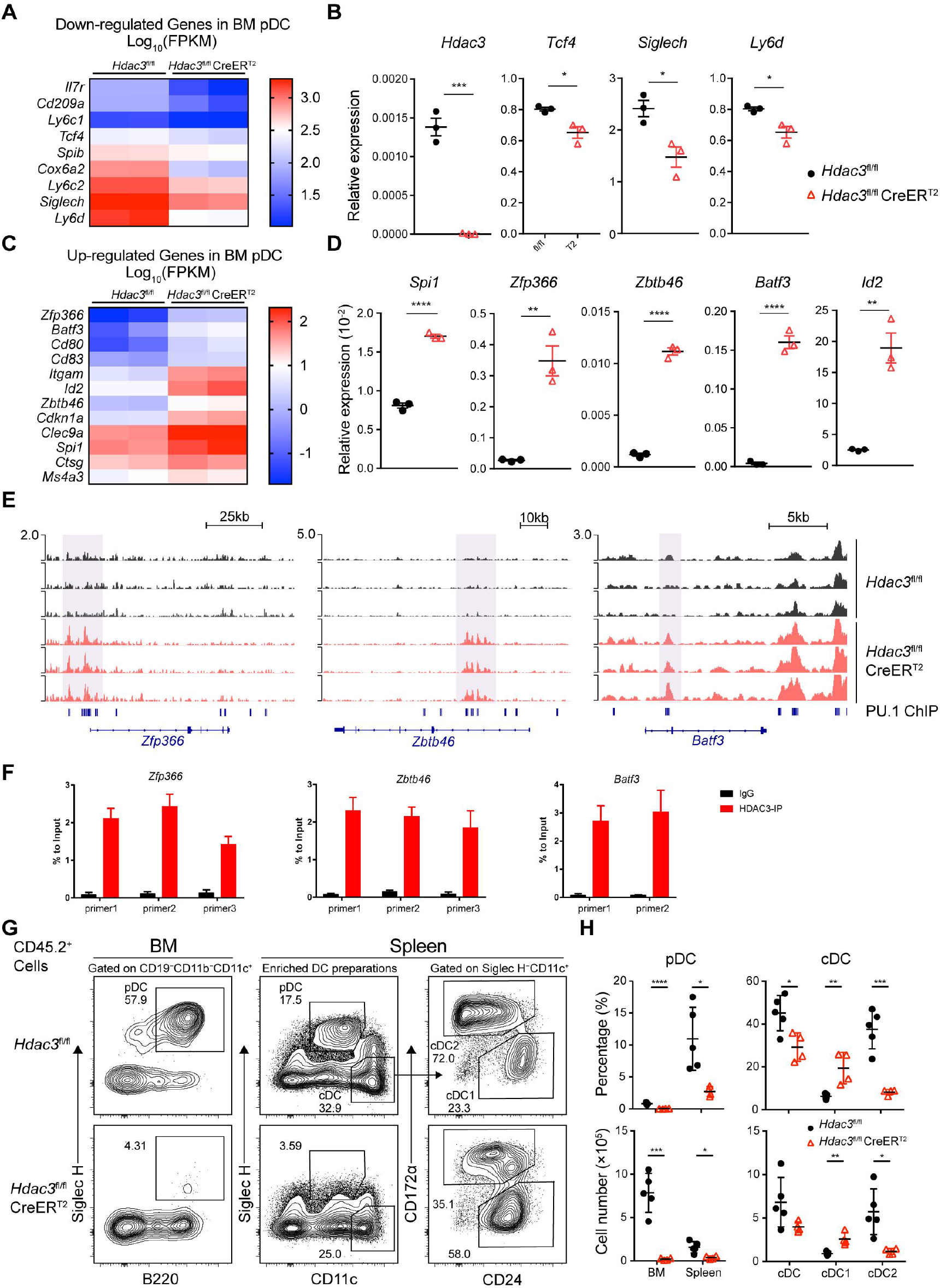
HDAC3 repressed H3K27ac around PU.1-binding sites at cDC genes loci. **(A-D)** Up-regulated and down-regulated DEGs (Fold Change > 2, q-value ≤ 0.05) in *Hdac3*-deficient BM pDCs compared with WT pDCs, determined by RNA-seq analysis (A, C) and confirmed by qRT-PCR (B, D). **(E)** Sequencing tracks of CUT&Tag analysis, with anti-H3K27ac antibody, of BM pDC sorted from *Hdac3*^fl/fl^ and *Hdac3*^fl/fl^ CreER^T2^ mice. Loci of cDC transcription factor *Zfp366, Zbtb46, Batf3* are displayed. **(F)** ChIP-qPCR validation of HDAC3 binding sites in BM pDCs on *Zfp366, Zbtb46, Batf3* loci identified in CUT&Tag analysis. Representative data of two independent experiments with similar pattern. **(G-H)** Lethally irradiated CD45.1 WT mice were reconstituted with BM alone from CD45.2 *Hdac3*^fl/fl^ or *Hdac3*^fl/fl^ CreER^T2^ mice. Eight weeks after reconstitution, *Hdac3* deletion was induced by Tamoxifen and BM cells and splenocytes were analyzed after three weeks. Representative flow cytometry profiles of CD45.2^+^ BM pDCs and splenic DC subsets (G), the percentage of DC subsets among CD45.2^+^ cells and absolute cell number (H) in BM chimeric mice. The staining profiles of pDC and cDC subsets on gated CD45.2^+^CD11b^−^CD19^−^CD11c^+^ cells from the BM and CD45.2^+^ enriched DC preparations from the spleens are shown. Results are from one experiment representative of 3 independent experiments with 4-5 animals each group. Data are shown as mean ± SD. * *p* < 0.05; ** *p* < 0.01; *** *p* < 0.001; **** *p* < 0.0001, by two-tailed Student’s *t*-test.

HDAC3 associates with NCoR/SMRT to form co-repressor complex and represses gene expression by suppressing histone acetylation modifications. To examine whether the up-regulated genes in HDAC3-deficient pDCs were repressed by HDAC3, we explored H3K27ac level in BM pDCs from *Hdac3*^fl/fl^ and *Hdac3*^fl/fl^ CreER^T2^ mice using CUT&Tag analysis. Motif analysis of unique H3K27ac peaks in HDAC3-deficient pDCs showed that these regions were significantly enriched with motifs recognized by PU.1 (SPI1) (**Figure 7 - Figure supplement 1A and B**). PU.1 promotes cDC development while repress pDC development by activating *Zfp366* expression(Chopin et al., 2019). HDAC3-deficient pDCs showed enhanced H3K27ac level around PU.1-binding region at cDC1-associated gene loci, including transcription factor *Zfp366, Zbtb46, Batf3* and cDC surface molecule *CD80, CD83* gene loci (**Figure. 7E and Figure 7 - Figure supplement 1C**). The binding of HDAC3 on the gene loci of the transcription factors *Zfp366, Zbtb46, Batf3* were further confirmed by ChIP-qPCR analysis in BM pDCs. As shown in **Figure. 7F**, significantly higher enrichments of HDAC3 on the sites of *Zfp366, Zbtb46, Batf3* loci than that of IgG control were observed. Since systemic Tamoxifen-induced *Hdac3* deletion in *Hdac3*^fl/fl^ CreER^T2^ mice led to death within 8∼9 days, we generated BM chimeric mice by transplanting BM cells from untreated CD45.2 *Hdac3*^fl/fl^ or *Hdac3*^fl/fl^ CreER^T2^ mice into lethally irradiated CD45.1 WT recipient. Eight weeks after reconstitution, recipient mice were treated with Tamoxifen, and DC subsets from BM chimeric mice were analyzed 3 weeks later to test the long-term DC differentiation ability of HDAC3-deficient progenitors (**Figure 7 - Figure supplement 1D**). Analysis of CD45.2^+^ donor cells showed that pDC development from *Hdac3*^fl/fl^ CreER^T2^ BM cells were severely defective. Meanwhile, the percentage and absolute number of splenic cDC1 derived from *Hdac3*^fl/fl^ CreER^T2^ BM cells were increased, while that of cDC2 were decreased (**Figure. 7G and H**). Taken together, these results suggested that HDAC3 could repress cDC1-associated gene expression through histone deacetylation and thereby promoted the expression of pDC-associated genes and subsequent pDC development.

## Discussion

In this study, we identified HDAC3 as a key epigenetic regulator for pDC development, evidenced by the absence of pDCs in BM and spleens of the HDAC3-deficient mice. Further investigation revealed that HDAC3 deficiency resulted in disturbed homeostasis of BM hematopoietic progenitors with DC differentiation potential. Moreover, we showed that HDAC3 was required for pDC differentiation from all early progenitors with DC differentiation potential. Mechanistic study by RNA-seq analysis comparing the gene expression profiles of the major progenitor populations and BM pDCs from HDAC3-deficient and control mice revealed significant downregulations of pDC-associated gene expression and upregulations of cDC1-associated genes. To further clarify how HDAC3 regulated the expression of these genes, we performed CUT&Tag analysis and ChIP-qPCR analysis. The results suggested that HDAC3 regulated the acetylation of H3K27 at cDC1-associated gene loci, thereby repressed the expression of cDC1-associated genes and promoted the expression of pDC-associated genes and subsequent differentiation of the progenitors towards pDCs. Consistently, HDAC3 deficiency resulted in decreased expression of pDC-associated genes and profound defect in pDC development. Thus, our study revealed a novel epigenetic regulatory role of HDAC3 in pDC development.

The development of DCs from hematopoietic progenitors is precisely orchestrated. We observed that deletion of *Hdac3* in resulted in marked decrease in the numbers of CMPs, CLPs and CD115^+^CDPs, which partially accounted for the reduced pDC generation. Whereas the numbers of HDAC3-deficient LSKs and CD115^-^CDPs were increased. However, despite the increase in the numbers, their abilities to differentiate into pDCs were severely impaired. The reduction in the number of CLP and increase in the number of LSKs in HDAC3-deficient mice were in line with the results of the BrdU incorporation analysis that showed a reduced DNA replication in HDAC3-deficient CLPs and an increased DNA synthesis in HDAC3-deficient LSKs. The incorporation level of BrdU in HDAC3-deficient and control CD115^+^CDPs and CD115^-^CDPs were comparable, though the number of HDAC3-deficient CD115^+^CDPs was reduced and CD115^-^CDPs increased. Considering that total number of CDPs did not change significantly, and a decrease of *Csf1r* (coding CD115) expression could be observed in RNA-seq analysis of the HDAC3-deficient CDPs. These results suggested that the increase in the number of CD115^-^CDPs might be due to the downregulation of *Csf1r* expression in CD115^+^CDPs. Overall, these results suggested that HDAC3 regulated different hematopoietic progenitors with different regulator mechanisms.

Cell fate determination is a key process during cell development, in which transcription factors play important roles. Recent studies reported PU.1-DC-SCRIPT-IRF8 axis as an important transcriptional pathway that regulated the fate determination between pDC and cDC1. PU.1 can activate *Zfp366* (encodes DC-SCRIPT) expression and DC-SCRIPT enhanced the expression of *Irf8* and thereby promoted the development and function of cDC1(Zhang et al., 2021). Loss of DC-SCRIPT leads to defective differentiation of cDC1 and an enhanced development of pDC(Chopin et al., 2019). These studies demonstrated that cell fate determination of cDC1 versus pDC was regulated in a mutually restrictive manner. In line with this, we also observed in the BM chimeric mice that HDAC3-deficient BM cells generated decreased number of pDCs but increased number of cDC1 compared to that derived from wildtype BM 3 weeks after Tamoxifen treatment, suggesting that HDAC3 may be involved in the fate determination of pDCs versus cDC1. This was confirmed by RNA-seq and qRT-PCR analysis of the residual HDAC3-deficient BM pDCs that showed down-regulation of pDC-signature genes such as *Tcf4, Siglech* and up-regulation of cDC1-associated genes, including *Zfp366, Id2* and *Batf3*, indicating that HDAC3 may promote pDC differentiation by repressing the expression of cDC1-associated genes. Further Cut & Tag analysis indicated that HDAC3-KO resulted in enhanced H3K27ac on *Zfp366, Zbtb46* and *Batf3* loci, and ChIP-qPCR analysis verified the binding of HDAC3 on these sites. Taken together, our study defined an essential role of HDAC3 in the regulation of pDC versus cDC1 lineage determination and revealed a new epigenetic regulatory mechanism for pDC development.

pDCs are the major producers of type I interferon and are involved in the development of some autoimmune diseases (Reizis, 2019). In this study we also found that HDAC3 specific inhibitor RGFP966 could efficiently block pDC generation *in vitro*, it therefore might serve as a potential therapeutic agent for pDC related diseases. Further investigation is warrant to evaluate this potential. Taken together, our study defined an essential role of HDAC3 in the regulation of pDC versus cDC lineage differentiation, and revealed a new epigenetic regulatory mechanism for pDC development.

## Materials and Methods

### Mice

All mice used in this study were C57BL/6 background. B6N-Tyr^c-Brd^*Hdac3*^tm1a(EUCOMM)Wtsi^/Wtsi mice were gifts from Prof. Fang-Lin Sun (Tongji University), FLP-DELETER mice: B6.129S4-*Gt(ROSA)26Sor*^*tm1(FLP1)Dym*^/RainJ (gain from Biocytogen, Beijing, China). *Hdac3*^tm1a(EUCOMM)Wtsi^/Wtsi were crossed with FLP-DELETER mice to remove the neomycin (Neo) cassette flanked by two FRT sites, thus generating *Hdac3*^flox/flox^ (*Hdac3*^fl/fl^) with *Hdac3* exon 3 floxed by a pair of loxP site. *Itgax*-Cre mice were gifts from Prof. Nan Shen (Shanghai Jiao Tong University) and B6.129-*Gt(ROSA)26Sor*^*tm1(cre/ERT2)Tyj*^/J (CreER^T2^) mice were purchased from The Jackson Laboratory. *Hdac3*^fl/fl^ mice were crossed with *Itgax*-Cre or CreER^T2^ mice to generate *Hdac3*^fl/fl^ *Itgax*-Cre mice and *Hdac3*^fl/fl^ CreER^T2^ mice. Mice were bred and maintained in a specific pathogen-free (SPF) animal facility at Tsinghua University, with 12/12-hour light/dark cycle, at 22-26 °C and sterile pellet food and water and libitum. All animal procedures were performed in strict accordance with the recommendations and approval of the Institutional Animal Care and Use Committee at Tsinghua University.

### Cell isolation and flow cytometry

For flow cytometry, BM cells and spleens were collected. Spleens were then minced into fine pieces and digested with Collagenase III (1 mg/ml, Worthington) and DNase I (0.1 mg/ml, Roche) in RPMI-1640 for 30 minutes at room temperature with rotating. Erythrocytes were removed by using Red Cell Removal Buffer (0.168M NH4Cl). Cell suspensions were filtrated through a 70 μm cell strainer. BM cells or splenocytes were then separated by density gradient centrifugation over 1.086g/mL or 1.077g/mL Nycodenz respectively (Sigma-Aldrich). The cell fraction harvested from the interface between fetal bovine serum (FBS) and Nycodenz was then incubated with antibody cocktail (BM Lin^−^ cells: anti-CD2, anti-CD3, anti-CD8α, anti-B220, anti-CD19, anti-TER119, anti-CD90, anti-CD11b, and anti-Ly6G; DC preparations from the spleens: anti-CD3, anti-CD19, anti-TER119, anti-CD90 and anti-Ly6G). Cells were washed and incubated with BioMag Goat Anti-Rat IgG (Bangs Laboratories) on ice. Beads were then removed using magnetic separator. Cell suspensions were incubated with homemade anti-CD16/32 antibody or Rat Gamma Globulin (Jackson ImmunoResearch) for blocking and then stained with fluorochrome-conjugated monoclonal antibodies against CD45, CD11c, Siglec-H, B220, PDCA-1, CD45RA, Ly6C, CD8α, CD172α, XCR1, and CD24. Cells were analyzed on the LSR Fortessa instrument (Becton Dickinson), and data were analyzed with FlowJo X software (TreeStar). Cell sorting was performed with a FACS Aria III cell sorter (Becton Dickinson) for subsequent experiments. Antibodies are listed in reagents session..

### FLT3L stimulated bone marrow culture

DC generation by FLT3L stimulated BM cultures were performed based on the original protocol reported by Naik et.al(Naik et al., 2005). Briefly, total BM cells were plated at 1.5×10^6^ cells/ml in RPMI-1640 complete medium, in the presence of 200ng/ml FLT3L (PeproTech) and 1μM 4-Hydroxytamoxifen (Sigma-Aldrich). Cells were harvested and analyzed on day 5-9. To generate DCs from progenitors *in vitro*, the progenitor populations LMPPs (10^3^), Flt3^+^ CMPs, CLPs, and CDPs (10^4^), were isolated and cocultured with 1.5 × 10^4^ CD45.1 BM feeder cells in the presence of 200ng/ml FLT3L with 1 μM 4-Hydroxytamoxifen. Cells were harvested and analyzed on day 4 (for CLPs or CDPs), day 5 (for CMPs) or day 9 (for LMPPs).

### Bone marrow reconstitution

To generate mixed BM chimeras, 1×10^6^ CD45.2 *Hdac3*^fl/fl^ or *Hdac3*^fl/fl^ CreER^T2^ bone marrow (BM) cells were mixed with CD45.1 wild type (WT) competitor BM cells at 1:1 ratio, and then transplanted into lethally irradiated CD45.1 WT mice. Eight weeks later, *Hdac3* deletion was induced by Tamoxifen treatment. To generate DCs from progenitors *in vivo*, 10^4^ LSKs or CD115^−^CDPs were sorted from *Hdac3*^fl/fl^ or *Hdac3*^fl/fl^ CreER^T2^ mice after Tamoxifen treatment, and then were transplanted into lethally irradiated CD45.1 WT mice mixed with 2× 10^5^ CD45.1 BM cells. Three weeks (for LSKs) or 10 days (for CD115^−^CDPs) post reconstitution, BM cells and enriched DC preparations were analyzed. To examine DC differentiation potential of HDAC3-deficient progenitors in the long-term, lethally irradiated CD45.1 WT mice were reconstituted with untreated BM of CD45.2 *Hdac3*^fl/fl^ or *Hdac3*^fl/fl^ CreER^T2^ mice. Eight weeks later, *Hdac3* deletion was induced by Tamoxifen and BM cells and splenocytes were analyzed three weeks post treatment.

### Retrovirus Production and Infection

Retroviral supernatants with pMYS-iresGFP, pMYS-HDAC3(WT)-iresGFP, pMYS-HDAC3(Y298F)-iresGFP were centrifuged onto RetroNectin (Takara)-coated plates for 2 hours at 2000 g at 32°C. Sorted BM Lin-cells were cultivated with the virus in the presence of IL-3, IL-6 and SCF for 48 hours. GFP+ cells were then sorted.

### Cleavage under targets and tagmentation (Cut & Tag) analysis

The Hyperactive In-Situ ChIP Library Prep Kit was purchased from Vazyme, and libraries were generated following the manufacturer’s instructions. Briefly, 1 × 10^5^ BM pDCs sorted from *Hdac3*^fl/fl^ control or *Hdac3*^fl/fl^ CreER^T2^ littermates were bound to beads and were subjected for immunoprecipitation with primary anti-H3K27ac antibody (Active Motif) at 4°C overnight. A secondary anti-rabbit antibody (Abcam) was then added and incubated under gentle agitation at room temperature (RT) for 1 hour. Cells were then washed and incubated in Dig-300 buffer with 0.04μM hyperactive pG-Tn5 transposon at RT for 1 hour. Cells were washed and then incubated in 100 μl of tagmentation buffer (10 mM MgCl2 in Dig-300 buffer) at 37°C for 1 hour. Reaction was stopped by addition of SDS, EDTA and proteinase K and incubated at 55°C for 1 hour and then DNA were extracted. For library amplification, 24 μL of DNA was mixed with 10 μL 5× TruePrep Amplify Buffer (TAB), 1 μL TAE, and 5 μL barcoded i5 and i7 primers and amplified for 20 cycles. PCR products were purified with VANTS DNA Clean beads (Vazyme). Libraries were sequenced on an Illumina NovaSeq platform.

### RNA extraction and quantitative RT-PCR

RNA of sorted cells was extracted by TRIzol Reagent (Invitrogen), and cDNA was reversely transcribed using PrimeScript^™^ RT Master Mix (Takara). For quantitative RT-PCR, PowerUp^™^ SYBR^™^ Green Master Mix (Applied Biosystems) was used, and samples were run on an HT7900 (Applied Biosystems) quantitative PCR machine. Gene expression was normalized to the expression of house-keeping gene Actin. Primers used: *Hdac3*: (forward) 5’-TGATCGTCTTCAAGCCTTAC-3’ (reverse) 5’-TTGGTGAAACCCTGCATAT-3’, *Batf3*: (forward) 5’-GGAACCAGCCGCAGAG-3’, (reverse) 5’-GCGCAGCACAGAGTTCTC-3’, *Spi1*: (forward) 5’-ATGTTACAGGCGTGCAAAATGG-3’, (reverse) 5’-TGATCGCTATGGCTTTCTCCA-3’, Zfp366 (forward) 5’-CTTCCTGCCCAAGCAGCCCC-3’ (reverse) 5’-GGCACTGCCAGCGCTTCTGA-3’, *Id2* (forward) 5’-ACCAGAGACCTGGACAGAAC-3’ (reverse) 5’-AAGCTCAGAAGGGAATTCAG-3’, *Zbtb46*: (forward) 5’-GACACATGCGCTCACATACTG-3’ (reverse) 5’-TGCACACGTACTTCTTGTCCT-3’, *Tcf4*: (forward) 5’-CGAAAAGTTCCTCCGGGTTTG-3’ (reverse) 5’-CGTAGCCGGGCTGATTCAT-3’, *Siglech*: (forward) 5’-GGAACCAACCTCACCTGTCA-3’ (reverse) 5’-AGAGACATGGGCTGTGGAGT-3’, *Ly6d*: (forward) 5’-CCTCAGCCTGCTCACTGTTA-3’ (reverse) 5’-CAAGGGAAATTCCAAGCAGT-3’, *Actin* : (forward) 5’-ATGCTCCCCGGGCTGTAT-3’ (reverse) 5’-CATAGGAGTCCTTCTGACCCATTC-3’

### Protein extraction and Western blot analysis

Sorted DC subsets were lysed in cold RIPA Lysis Buffer (Beyotime Biotechnology) supplemented with protease inhibitor cocktail (Roche), phosphatase inhibitor cocktail (Bimake), and SDS loading buffer (Beyotime Biotechnology). Total protein was denatured and loaded into the SDS-PAGE, and then transferred to PVDF membranes. After blocking, primary antibodies were hybridized overnight (HDAC3, 10255-1-AP, ProteinTech; β-actin, 3700, CST) and then secondary antibodies. Imaging was carried out with Amersham Imager 600 System (GE).

### *In vivo* expansion of DCs with Flt3L-secreting B16F10 melanoma

To acquire abundant pDCs for ChIP assays, we utilized a Flt3L-secreting B16F10 melanoma cell line (B16F10-Flt3L) to stimulate DC expansion *in vivo*. B16F10-Flt3L cells were expanded in complete RPMI 1640 medium. Upon inoculation, 2 × 10^6^ cells were subcutaneously injected to the nape of the neck of each wild-type male C57BL/6 mouse. The mice were monitored and sacrificed for BM pDC purification as previously described 14 days post inoculation.

### ChIP-qPCR analysis of HDAC3 binding sites in BM pDCs

ChIP-qPCR analysis were performed using formaldehyde fixed purified BM pDCs with ChIP-IT^®^ Express Chromatin Immunoprecipitation Kits (53035, Active Motif, USA) according to the manufacturer’s instructions. Briefly, sorted BM pDCs were fixed using 1% formaldehyde for 10 min and quenched with 10 × glycine in the kit. Then cells were washed with PBS for 3 times and resuspended using ChIP lysis in the kit. After centrifuged at 2000rpm for 5 min, the cells were the lyzed in digest buffer with proteinase inhibitor cocktail and phosSTOP (Roche, Switzerland) on ice. Then the cells were sonicated using Bioruptor® Pico sonication device (Diagenode, Belgium) with 30s on/ 30s off for 5-6 cycles at 4°C, and then add enzymatic shearing cocktail to the system and incubated at 37°C for 8 min, reaction was stopped by adding of EDTA and the cell lysate was centrifuged at 12000rpm for 10 min. Before diluted in ChIP dilution buffer (with proteinase inhibitor cocktail and phosSTOP) appropriate volume of supernatant was collected as 20% Input. Then the sample were divided into two equal volumes and antibody of HDAC3 (85057S, CST, USA) or normal rabbit IgG (2729S, CST, USA) was added according to the antibody instruction and incubated at 4°C overnight. Then 10 μl of pre-washed Magna ChIP^™^ Protein A+G Magnetic Beads (16-663, Millipore, USA) were added to each sample respectively and incubated for 2.5 h. Then the beads were washed sequentially with low salt buffer, high salt buffer and TE buffer at 4°C, the beads were then washed by TE buffer again and then remove the entire supernatant. Then the beads sample and input were added 172 μl TE buffer, 10 μl 10% SDS and 8 μl 5M NaCl, and incubated at 65°C overnight shaking vigorously. RNase A in the kit were then added and incubated at 37°C for 15 - 30 min followed by adding the proteinase K (20 mg/ml) to the system and incubated at 55 for 1 h. The beads were then removed and DNA fragments in the supernatants were extracted using MinElute PCR Purification Kit (28004, Qiagen).

Quantitive Real time PCR were performed with primers:

*Zfp366*-primer1: (forward) 5’-CTTTGGATCGGGACTGGACC-3’,

(reverse) 5’-TGAATGGGGAGCCATAGGGA-3’;

*Zfp366*-primer2: (forward) 5’-GTGCTGGGGTTTCAAAGCAG-3’,

(reverse) 5’-GGGCTAACCAAGAGGGAACC-3’;

*Zfp366*-primer3: (forward) 5’-GCCACCACAAAGAACCACAC-3’,

(reverse) 5’-GAGCTGGGCCCAAATCATCT-3’;

*Zbtb46* primer1: (forward) 5’-GAGGAACAAGGTAGCCCCAG-3’,

(reverse) 5’-CCCATACACCACTTGCCCTT-3’;

*Zbtb46* primer2: (forward) 5’-TCACATCTGGGTGGGATTGC-3’,

(reverse) 5’-TCCGTTGCTGTCACGGTTTA-3’;

*Zbtb46* primer3: (forward) 5’-CTTGCCTAGCACCCAGCTAA-3’,

(reverse) 5’-GCAAACCCATCCAATGCTCC-3’;

*Batf3* primer1: (forward) 5’-TGCACAGCAAGTTCTAGCGA-3’,

(reverse) 5’-TCCCCAAACCAACGTTCACA-3’;

*Batf3* primer2: (forward) 5’-GGGGCAGAAGTTTGTGAACG-3’,

(reverse) 5’-GTCAGGCCCTGTGTTCCATA-3’.

### Statistical analysis

Analysis of all data was done with unpaired two-tailed Student’s *t*-test (Prism, GraphPad Software). *p* < 0.05 was considered significant. \**p* < 0.05; \*\**p* < 0.01; \*\*\**p* < 0.001; \*\*\*\**p* < 0.0001

### Reagents

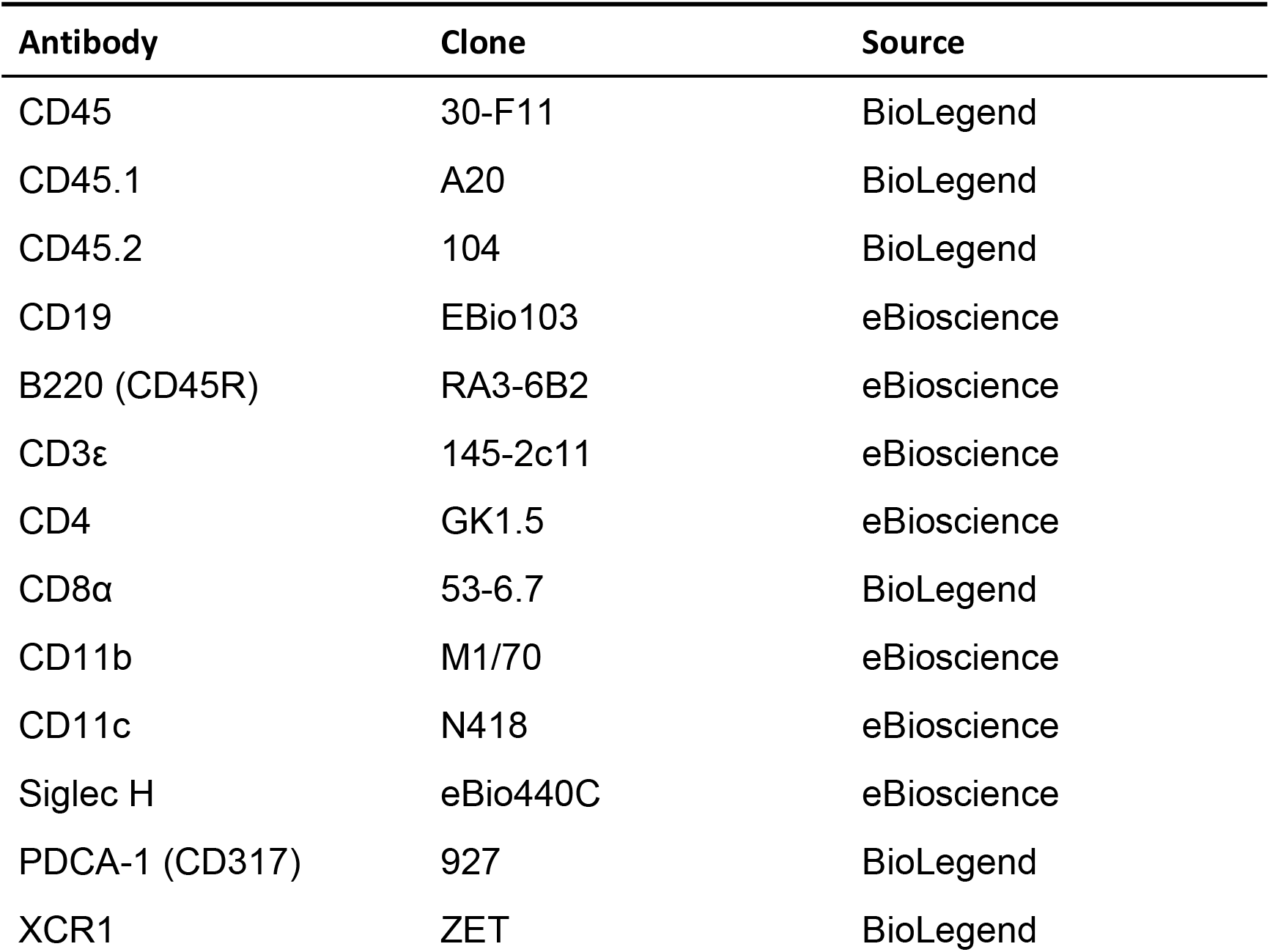
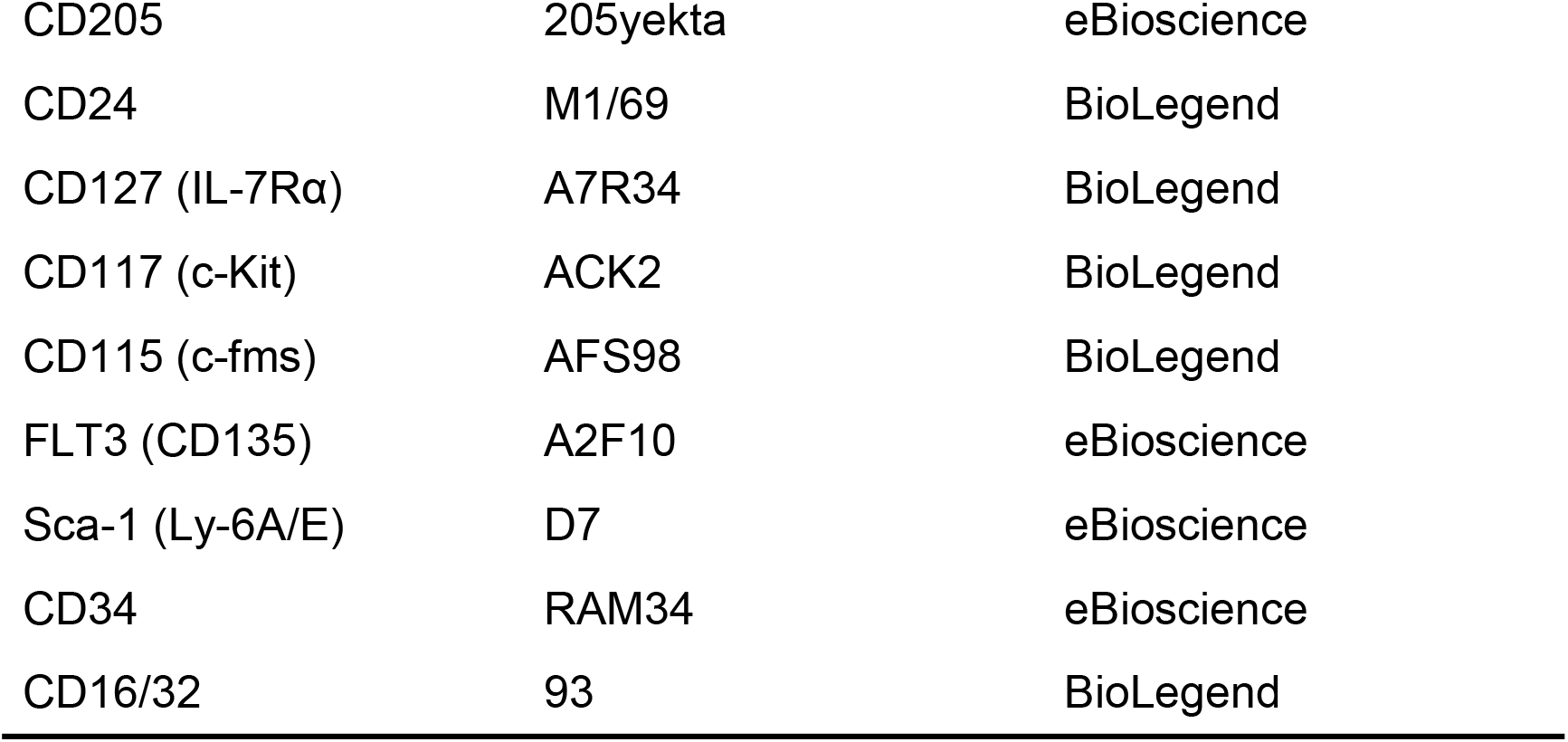

| Antibody | Clone | Source |
| --- | --- | --- |
| CD45 | 30-F11 | BioLegend |
| CD45.1 | A20 | BioLegend |
| CD45.2 | 104 | BioLegend |
| CD19 | EBio103 | eBioscience |
| B220 (CD45R) | RA3-6B2 | eBioscience |
| CD3ε | 145-2c11 | eBioscience |
| CD4 | GK1.5 | eBioscience |
| CD8α | 53-6.7 | BioLegend |
| CD11b | M1/70 | eBioscience |
| CD11c | N418 | eBioscience |
| Siglec H | eBio440C | eBioscience |
| PDCA-1 (CD317) | 927 | BioLegend |
| XCR1 | ZET | BioLegend |
| CD205 | 205yekta | eBioscience |
| CD24 | M1/69 | BioLegend |
| CD127 (IL-7Rα) | A7R34 | BioLegend |
| CD117 (c-Kit) | ACK2 | BioLegend |
| CD115 (c-fms) | AFS98 | BioLegend |
| FLT3 (CD135) | A2F10 | eBioscience |
| Sca-1 (Ly-6A/E) | D7 | eBioscience |
| CD34 | RAM34 | eBioscience |
| CD16/32 | 93 | BioLegend |

### Data availability

For original data, please contact

RNA sequencing data are available at NCBI GEO Datasets under accession GSE197207, https://www.ncbi.nlm.nih.gov/geo/query/acc.cgi?acc=GSE197207

Cut & Tag data are available at NCBI GEO Datasets under accession GSE197212, https://www.ncbi.nlm.nih.gov/geo/query/acc.cgi?acc=GSE197212

## Author contributions

L.W. and Y.Z. designed the research, Y. Z. and Z.H. performed experiments and analyzed the data, W. L. generated HDAC3-deficient mice. X. S. assisted with performing some experiments. T. W. and J. L. assisted with analyzing the data and editing the manuscript. Y. Z., Z. H. and L.W. wrote the manuscript. L.W. supervised the research.

## Acknowledgments

We thank Prof. Fang-Lin Sun (Tongji University, China) for providing B6N-Tyr^c-Brd^*Hdac3*^tm1a(EUCOMM)Wtsi^/Wtsi mice and Prof. Nan Shen (Shanghai Jiao Tong University, China) for providing *Itgax*-Cre mice. We are grateful for the support provided by the animal core facility at Tsinghua University. This research was supported by the Ministry of Science and Technology of China (National Key Research Project 2019YFA0508502 to L. Wu), the National Natural Science Foundation of China (grants 31991174 to L. Wu, 31800769 to Z. He), Funding from Tsinghua-Peking Center for Life Sciences (to L. Wu).

